# Epilepsy in children in Papua New Guinea: a longitudinal cohort study

**DOI:** 10.1101/551416

**Authors:** Casparia Mond, John D Vince, Trevor Duke

## Abstract

**Background:** Epilepsy affects up to 1-4% of children living in low income and middle countries, however there are few studies of the problems faced by children with epilepsy in such settings.

**Aim:** To document the characteristics and situation for children with epilepsy in Port Moresby, an urban area in Papua New Guinea, a low-middle income country in the Western Pacific region. To describe the types of epilepsy, associated comorbidities, treatment access and barriers, the adequacy of seizure control, the quality of life and developmental opportunities, and the difficulties faced by children with epilepsy and their families.

**Methods:** A longitudinal cohort study, following children with epilepsy over 18-24 months. Mixed methods evaluations included assessments of seizure control, medications, neurodevelopment, and structured interviews with children and parents, and a parent-diary to record additional information.

**Results:** Forty-seven children with epilepsy were followed for a median of 18 months; 75% were being treated with phenobarbitone. Seizure control improved over time for some children, but inconsistent supply of phenobarbitone hindered better control. Twenty six (55%) children had some developmental delay. Children gave vivid descriptions of their experience of seizures. Most children and parents had a positive view of the future but faced many challenges including financial difficulties, fear of seizures especially at school, restriction of activity and stigma and discrimination.

**Conclusion:** Comprehensive care for children with epilepsy requires a good knowledge of the individual patient - including their seizure type and comorbidities, their family, and their strengths and vulnerabilities. It requires long term follow up, with a dedicated team of health professionals to provide support.

## Introduction

Epilepsy is the most common neurological disorder in children with prevalence estimates in low income countries varying from 3.6-44 per 1000 in children.(1-3) WHO estimates that 80% of affected children live in countries with limited resources, and the treatment gap where affected children are not consistently treated is estimated at greater than 60%.(4-6) The consequences of inadequate treatment are often severe. These include preventable seizures and secondary injury from falls, burns and drowning.(5) Furthermore, many children with epilepsy suffer loss of potential for education and development, and have sub-optimal school and community participation. There are significant mental health consequences of having poorly controlled seizures, including anxiety and fear, isolation from peers, and feelings of stigma and discrimination for affected children and their families.(7) Poor control of seizures or inadequate treatment of people with epilepsy increases the risk for sudden unexpected death in epilepsy (SUDEP), a rare but poorly understood complication.(8)

Little is known about epilepsy and its comorbidities in children in Papua New Guinea (PNG) or Pacific Island communities. There has been only one previous study from PNG - a prospective study of 40 children at Port Moresby General Hospital in 1990-91.(9) Our aim was to follow a cohort of children with epilepsy and document its nature and extent, associated comorbidities, access to treatment, the adequacy of seizure control, the quality of life and developmental opportunities, and the difficulties faced by them and their families.

## Methodology

Children between the ages of 1 and 18 years with epilepsy who attended Port Moresby General Hospital (PMGH) were included in this study. Children were recruited from the outpatients department, the children’s wards, the emergency department and the neurology clinic. Children entered the study after their parents gave written consent. The parents were interviewed using a structured questionnaire with closed and open questions. Children who were old enough and had the communication skills to answer questions were also interviewed if their parents consented.

The questionnaire was designed to obtain baseline demographic data, a description of the convulsions to enable classification the AEDs given to the child, and issues associated with access to and effectiveness of AEDs, and additional medical care required by the child. We sought to understand parental concerns and challenges faced in caring for their child, and the extent and challenges of school participation.

The children’s questionnaire contained questions about the child’s perception of his or her illness and why the AED was taken, as well as questions about stigma or discrimination they may have experienced, and their hopes for their future.

Interviews were conducted in English or Tok Pisin as appropriate. Some interviews were recorded electronically and were later transcribed if consent was given. A neurodevelopment assessment included motor, visual, hearing, speech and language assessment using the Denver II Developmental Assessment for children aged 1 year to 6 years. For older children a neurological examination was done and neurological impairments documented.

The parents or guardians of 28 children were given a diary. Those not given diaries were either illiterate or felt they would not use the diary as their child did not have frequent seizures. The diary was used to record the number, type and duration of seizures, any aura the child may have experienced, if anticonvulsants were missed and the reasons why. Problems with school attendance and any other problem or questions the parents might wish to ask the doctor at the next review were also to be noted. These diaries were reviewed at each follow up session and the content noted.

The degree of seizure control was classified as good (1-4 seizures per month); moderately well controlled (5-10 seizures per month) or poorly controlled seizures (11-30 seizures per month).

### Follow-up

Reviews occurred monthly if the seizures were poorly controlled, or every 2-3 months for other children, depending on parents’ availability and the distance they lived from the hospital.

Data were entered into an Excel spread sheet and analysed descriptively. Qualitative data are presented as narrative or thematically.

### Ethics approval

Ethics approval was given by the University of Papua New Guinea School of Medicine and Health Science Research Committee, and written informed consent was gained from parents of all children involved.

## Results

The cohort consisted of 47 children with epilepsy, of which 25 were female. Table 1 describes the characteristics of these children. The median age was 6.5 years (interquartile range 5.5–12 years). Twenty-one children had normal development for their age. Eleven had gross motor dysfunction and 14 had delayed fine motor skills. Four had visual impairment and two had hearing impairment. Nine had global developmental delay.

**Table 1.** Patient characteristics and comorbidities.

|  |  |
| --- | --- |
| **Patients (n=47)** |  |
| Median age (years) (IQR) | 6.5 (5.5 - 12) |
| Male: female | 22:25 |
| Normal development for age (%) | 21 (45) |
| **Comorbidities** | **No. (%)** |
| Gross motor dysfunction | 20 (42) |
| • Hypotonia | 1 (2) |
| • Hypertonia | 3 (6) |
| • Hemi or monoplegia | 11 (23) |
| • Spastic quadriparesis | 5 (11) |
| Fine motor skills delay | 14 (30) |
| Vision impairment | 4 (9) |
| Speech impairment | 16 (34) |
| Hearing impairment | 2 (4) |
| Behaviour problems or neurodevelopmental comorbidities | 18 (38) |
| • Hyperactive | 11 (23) |
| • Mental health / psychosis | 4 (9) |
| • Poorly responsive | 3 (6) |
| Global developmental delay | 9 (19) |
| Difficulty feeding | 6 (13) |

Table 2 documents the seizure types and AEDs used by the children at baseline. The majority (70%), were taking phenobarbitone monotherapy.

**Table 2.** Seizure types and anti-epileptic drugs.

| **Types of seizures** | **Number of patients**<br>**n=47 (%)** |
| --- | --- |
| Generalised tonic-clonic | 24 (51) |
| Tonic | 3 (6) |
| Myoclonic | 2 (4) |
| Partial simple | 2 (4) |
| Atonic | 1 (2) |
| Absence | 1 (2) |
| Partial complex | 0 (0) |
| Combination of seizure types | 14 (31) |
| **Antiepileptic drugs** |  |
| Phenobarbitone monotherapy | 33 (70) |
| Phenytoin monotherapy | 1 (2) |
| Sodium valproate monotherapy | 1 (2) |
| Diazepam monotherapy | 1 (2) |
| Carbamazepine monotherapy | 0 (0) |
| More than one AED: mostly phenobarbitone and carbamazepine | 11 (23) |

Table 3 documents the severity of the epilepsy and seizure control at baseline and follow up. At recruitment, 35 (74%) children were classified as having good seizure control 5 (11%) had moderately well controlled seizures. Two children had poorly controlled seizures and 5 (115) children were considered to have very poor control having more than 30 seizures a month.

**Table 3.** Seizure control at baseline and follow-up.

| Seizures/month | Baseline | 5 months | 10 months | 15 months | 20 months |
| --- | --- | --- | --- | --- | --- |
|  | n =47 (%) | n = 47 (%) | n = 42 (%) | n = 43 (%) | n= 23 (%) |
| Good <1-4 | 35 (74) | 36 (77) | 35 (83) | 23 (88) | 21 (92) |
| Moderate 5-10 | 5 (11) | 5 (11) | 3 (7) | 1 (4) | 1 (4) |
| Poor 11-30 | 2 (4) | 4 (8) | 4 (10) | 2(8) | 1 (4) |
| Very poor >30 | 5 (10) | 2 (4) | 0 | 0 | 0 |

Five children had sustained injuries during seizures, including a flame burn of the arm and facial injuries and several had scars from lacerations from falls, and from tongue and finger biting.

Only 3 of the 28 parents who were given diaries used them diaries and brought them regularly to the clinic. One of these were the parents of a child with very poorly controlled seizures. The diary contents was checked monthly and AEDs were adjusted accordingly. Another parent found the diary useful enough to keep using it consistently throughout the follow-up period. She recorded her child’s seizures, visits to the physiotherapist, the problems they faced and her questions. AEDs were also adjusted based on the diary notes.

### School attendance

Only nine (26%) of the 35 school-aged children were attending primary school at the time of study recruitment. None were in high school. One had completed primary school (Grade 8 at age 16 years) two years prior to this study and was at home planning to start a small business. Sixteen children had been enrolled at primary or elementary schools in the past but had ceased attending because they had a seizure at school. Many parents feared further seizures in school with no family member to help the child. Some children had been withdrawn from school because of hyperactivity or being unable to concentrate in class.

### Children’s interviews

17 children between 7 and 18 years were interviewed. The panel lists the questions asked and some of the verbatim answers. Most of the children were able to give a description of their seizures or what they felt at the onset of seizures. For example, five (29%) children) said they had headaches or felt heaviness of the head, and three children described blurred vision and dizziness. Some gave descriptions such as “teeth locked” or “legs stiff”. Children were also asked if they knew what epilepsy was and over half of them responded that they had some sickness in the head and would shake (“guria guria”).

The children were asked what they liked or did not like about school. Of the 17 children interviewed 9 were attending school at the time they were interviewed and 5 had been to school previously, and 3 had never been to school. They nominated the following reasons for enjoying school: because they had friends at school, because they liked certain subjects at school such as English, science, reading and writing, one child liked playing sports at school, and two liked going to school because they liked their teachers. A 7 year old in first grade stated that he liked going to school because his teacher let him sharpen his pencils with the large sharpener in his classroom.

Questions about whether they were treated differently by their family, friends or teachers were answered by 15 children. Two children just smiled when asked but did not reply. 14 of the 15 children (93%) who answered said they were treated differently by their friends and family because they had epilepsy. They all had some restrictions placed on them by their parents so they could not go out and play as often as their siblings and were restricted from spending time away from their homes such as spending a weekend away at a friend or relative’s house.

All the children expressed their wish to have few or no restrictions placed on them. They did not like to be teased. Only one child mentioned that the restrictions were for his own good so he accepted them although he was unhappy at times as a result.

In response to the question ‘What would you like to be when you are older?’ most of the children were enthusiastic about the future. Twelve children knew what they wanted to do in the future while four said they did not know and three children did not answer the question. Five children wanted to be doctors while other choices were a pastor, a lawyer, a pilot, a fisherman. One wanted to work with computers and one wanted to work in an office and write in books.

Children were asked about any worries or concerns they may have. Sixteen children answered this question. Worries and concerns were mostly related to epilepsy, their family or school. Five children had worries directly related to epilepsy, wanting their seizures to stop for good.

### Parents’ interviews

The 47 children in this study had either one or both parents or a guardian who participated in answering the parents’ questionnaire (see Panel for indicative verbatim responses). When asked the question ‘What do you know about epilepsy?’ the parents gave various answers based on what part of the body they thought was affected or what they had read or been told by doctors or from their experiences of seeing their child’s seizures. Thirty (64%) parents answered that epilepsy was some sort of brain disorder. Six children (13%) had at least one relative who also has or had seizures in the past so their parents believed that epilepsy is an inherited condition. Another four parents (8.5%) answered that they were not sure what condition their child had and so were not sure what epilepsy was, and 4 believed that magic or the supernatural was involved.

Parents were asked whether they thought epilepsy was treatable or not. Thirty-three (70%) answered that epilepsy was treatable while four caregivers (9%) stated that epilepsy was not. Ten (21%) parents believed that epilepsy was sometimes but not always treatable.

The parents were also asked what challenges their children faced in attending school. Twenty-four parents whose children had at some point attended school answered this question. Fourteen (58%) said their child had no challenges in school and the child enjoyed going to school. A 14 year old enjoyed school although she was sometimes teased. Even when her family had no bus fare or lunch money for her and she was told to stay at home she would walk to school and not worry about the money. Ten (42%) parents stated that their child faced challenges in school in terms of difficulty learning due to poor memory, being hyperactive in class and being slow in reading or writing. Other parents stated that their children were teased in school and one child was not allowed to participate in school sports because he had epilepsy.

Forty-six parents answered the question on challenges they faced. The main challenges were financial, the constant care required, their child’s behaviours and stigma. Some parents expressed multiple challenges others did not find caring for their child a challenge.

Thirteen parents (28%) identified financial challenges as the main one. They found it difficult to buy AEDs when the hospital pharmacy ran out, and they also faced challenges with bus fare and care of the child especially when going for monthly clinic or physiotherapy visits. The cost of phenobarbitone in private pharmacies ranged from K36.00 – K210.00 per month (US$9-52). Parents of children with disabilities had higher financial challenges. A major challenge for some of the parents was always having someone to watch the child in case of a seizure or an accident. Nine parents (20%) mentioned this as their main challenge. Nine (20%) of parents reported their child’s behaviour, (hyperactivity, offensive language, tantrums, and difficulty following instructions) as their main challenge. Three parents had to leave work to stay home and watch their hyperactive child in case the child got hurt while playing or climbing trees or had a seizure while swimming as they lived near the sea.

Forty six parents gave their views of their child’s future. The majority were positive. Two-thirds stated that the child’s future would be fine or good “em bai orait” (he/she will be ok). Parents of 13 children (29%) were unsure of their child’s future and 2 parents (4%) of children with spastic quadriparesis stated that the future of their child was going to be poor. One grandmother stated sadly “she must go before me, that’s my prayer as there is nobody to care for her apart from me”.

Parents were also asked how the care of their child by the health services could be better. Issues raised were the long waiting time at the hospital pharmacy and the frequent stock-outs of phenobarbitone and other AEDs. Many suggestions were around communication and provision of information. Parents spoke of the need for doctors to better explain their child’s condition, and for doctors to spend time finding our more about the child’s condition and to ask about other issues affecting the child instead of just the AEDs and seizures.

A single mother who carries her 7 year old son (with quadriplegia) to physiotherapy weekly, suggested having social worker and physiotherapists at the neurology clinic as it was difficult taking hyperactive children or children who couldn’t walk and had to be carried to the social workers or physiotherapists on different days of the week.

### Follow-up

All 47 children were followed up for 15 months to 20 months depending on the date of recruitment into the study. Each child was seen monthly, every second or every third month depending on the parents availability and how far they lived from the hospital. There was a trend towards improved seizure control (73% vs 92% good control at 20 months, p=0.12).

## Panel

### Responses to questions from interviews with children with epilepsy, and their parents

If the response was in Tok Pisin, this is presented first, followed by the English translation

#### Children’s interviews

##### What do you feel when you get sick or have seizures?

- A 16 year old described his experience in this way “Mi sa feelim olsem mi no sa lukluk, hard lo lukluk, brain stap lo narapela hap, mi sa harim ol man toktok but sik controlim brain blo mi na mi hard lo toktok” (I feel like I cannot see, it’s hard to see, my brain is somewhere else, I can hear people talking but the sickness has controlled my brain so I cannot talk).
- One child described what she experiences as “Mi sa guria guria, tupla ear drum hot na brain sa move inside lo het” (I shake, my two ear drums are hot and my brain moves inside my head). Two children who had palpitations or mild chest pain prior to a seizure describe epilepsy as being related to the heart. One 9 year old said “lewa blo mi sa kisim shock” (my heart gets a shock).
- A 15 year old stated “em mas lek blo mi sik bikos em start olgeta taim lo lek na spread go lo het na mi blackout. Lek blo mi sa tait na mi sa sigaut bikos mi laik pudaun (it must be my leg that’s sick because it always starts with my leg and spreads to my head and then I black out. My legs becomes stiff and I scream because I am going to fall down.)

##### What do you think about school?

- A 17 year old who completed only one year of elementary school at age 8 years said “Mi laik go skul but mi no sa harim toktok blo teacher na ol mangi sa tok ‘longlong’” (I wanted to go to school but I never listened to what the teacher said and the kids would call me dumb)
- A 14 year old in fifth grade said “mi laik kisim save na mi bai wok na lukautim mama na papa” (I want to get educated so I can get a job and take care of my parents).

##### Are you treated differently by friends or family because of your epilepsy?

- One 9 year old who likes to play with his friends said ‘mama stopim mi lo go play na sa ok yu sikman ya stap lo haus, mi sa belhat. Ol sa givim mi moni na mi sa stap lo haus”. (Mum stops me from going to play and says you are a sick person so stay at home and I get angry. They (family) give me money so that I can stay home.) An 11 year old boy said “mum and dad stop me from playing and my brother calls me ‘sick dog’. My friends at school tease me and I hit them sometimes.”
- A 14 year old in grade 5 stated that her family do not allow her to go to her friends and relatives homes. Boys at school tease her and call her ‘guria guria meri’ (shake shake girl) and imitate her seizures as she has had several seizures at school since she started school. Another 16 year old in grade 8 is teased by kids in his village a lot and they call him ‘sik muruk’ (another term for seizure or epilepsy in Tok Pisin).
- One girl who was usually teased and isolated by her two younger sisters said “Ol sister blo mi mas lukautim mi na mipla play wantaim” (my sisters should take care of me and we play together).

##### What were their worries or concerns?

- Two children were concerned about school. One 16 year old who was in the last year of primary school (Grade 8) was worried she would not be allowed by her parents to go to high school as her classmates would not know what to do if she had a seizure in high school. Her friends at her current school know what to do and have helped her in the past when she had seizures at school. Another 15 year old was worried she could not trust her friends at school as they would go out with boys they knew she liked.
- The other 5 children had worries concerning their family. A 14 year old worried a lot about her deceased grandmother as her grandmother had help raise her and had passed away 3 years prior. An 11 year old boy missed his mother who lived in another province. A nine year old who lived with his mother was worried that his father did not care for him as his father had divorced his mother and started another family.

#### Parent’s interview

##### What do you know about epilepsy?

- Most parents stated that something was wrong with the brain itself “em sik lo brain” (it’s a sickness of the brain) or “em gat sik lo het” (he has a sickness of the head). Parents of four children thought it affected the brain because of worry or psychological problems in the child, three parents said it was post infection such as cerebral malaria or meningitis.
- One parent who thought epilepsy occurs post infection said “ol mosquito putim kiau lo head blo em taim em baby na em kisim malaria lo brain na bihain kisim displa sik” (mosquitoes laid their eggs in her brain when she was a baby and she got malaria in her brain then later got this sickness - epilepsy.
- Four parents believed that the origin of their child’s epilepsy is not medical but has a supernatural origin. One parent stated “ol man lo ples bagarapim em” (he has been cursed by people in our village).
- Three parents (6%) whose children always had generalized tonic clinic seizures believed that epilepsy was a condition which affected the whole body because their seizures affected their whole body. A mother of a 10 year old said “olgeta body blo em sa move taim em fit so sik so olgeta body em displa sik affectim” (the whole body moves when he has a fit therefore this sickness affects the whole body).

##### Main challenges faced by parents

- One parent of a 16 year old who has quadriplegia (and the fourth of five children) said “it’s (finances) a very big challenge. We have difficulty feeding her three meals a day as she is bigger now and eats a lot. We cannot afford transport to take her with the wheelchair for physiotherapy weekly”.
- Parents of 2 (4%) children felt that their main challenge was the stigma they perceived from the community. One parent of a 3 year old said “we don’t want people to say that we have a child with this sickness”.

##### How can health care for the child be better?

- One parent stated “doctors have to spend more time with the patient and parents and encourage them to put their children in school so parents won’t be afraid”. Another parent said “Doctors follow up in terms of medicine but finding out more about the patient will make the parents feel like they are concerned”. One parent whose child cannot speak said “pikinini blo mi ino sa toktok so dokta mas askim mi more lo wanem needs blo pikinini, ino raitim marasin tasol” (my child doesn’t speak so the doctor should ask me more about what the child’s needs are and not just write the medicine [prescription]).

## Discussion

There are few longitudinal studies of children with epilepsy in low and middle income countries.(10) This is one of the first studies in the Asia Pacific region to assess the adequacy of seizure control over time, and perceptions and aspects of quality of life for children with epilepsy and their families.(11) The follow-up period in our study averaged 18 months, sufficient to document outcomes, the challenges faced and the aspects of care that need to be considered in providing holistic services. The study also explored the children’s knowledge, ideas and understanding related to their epilepsy. These included the fears from their experience of seizures, their concerns regarding stigma and restrictions placed on their activity, the risk of injury during seizures, and the importance of their self-esteem in going to school, despite difficulties.

Whilst the majority of parents indicated that epilepsy was a brain disorder or an inherited problem, four answered that they were not sure of the cause, but another 4 said they thought some magic was involved. In studies in low and middle income countries belief that sorcery or witchcraft is a cause of epilepsy is often cited as a common cause of epilepsy.(12-14). This belief was not commonly voiced by the parents in our study, but it was volunteered by some, and may be more common in some less informed segments of their communities, and contribute to stigmatisation.

WHO recommends phenobarbitone as the first drug of choice for most seizures and epilepsies in developing countries because of its low cost and general effectiveness.(15) Phenobarbitone controls a range of seizure types and is on the essential drug list of 95% countries surveyed by WHO.(16) In our cohort seizure control was affected by the inconsistent supply of phenobarbitone at the public hospital pharmacy at Port Moresby General Hospital. Many parents could not afford to purchase phenobarbitone through private pharmacies. Phenytoin, carbamazepine and sodium valproate are also on the WHO essential drug list.(17) However, sodium valproate was not available at the public pharmacy and it was very costly to purchase elsewhere. Similar problems have been shown in other low resource settings.(18)

Nearly three-quarters of the children in this study had good seizure control based on the classification we used of 0-4 seizures per month. There are no standardised internationally accepted classification systems for assessing degree of control. Ours was an arbitrary and very basic classification. The number of seizures is not the only measure of adequacy of control, and duration of seizure and seizure type are important aspects to consider. There was a trend to improved seizure control over time, with fewer children having poor or very poor control of seizures after 18 months follow-up.

Previous studies of the use of seizure diaries in children, especially with multiple complex seizure types, identified limited reporting.(19) There are no studies that we could find of the use of seizure diaries in low resource settings. Compliance with use of the seizure diaries in our cohort was similarly limited; the diary being used regularly for only three children. The parents of these children found that keeping a diary was useful and their description of seizures and seizure frequency helped in their child’s management. The use of a diary requires a degree of literacy, and for families that are not literate some other way of recording the seizures could be devised.

Most children with epilepsy can go to school with support from their family, if their seizures are controlled and any associated complications are managed well. The parents in our study feared that seizures would occur at school and that the child would not cope in school due to distractibility or hyperactivity. Whilst some parents’ fears were based on past experience, even the parents of children whose seizures were adequately controlled feared putting their children in school. Having a simple plan written and discussed with the teacher and having emergency contacts at the school can help. A letter from the doctor informing the teacher of the child’s condition and what to do in the case of a seizure will also help. Despite parents’ feelings about their children’s challenges at school and some experience of stigma, almost all the children interviewed (15 out of 17) enjoyed attending school.

Some children volunteered vivid descriptions of their seizures and any aura they experienced. To our knowledge this has not been described in a systematic way before, and gives insights into children’s experiences and explains some perceptions and fears.

Many parents stated that they would like the doctors to better explain their child’s condition, spend time finding out more about the child’s condition and also discuss issues in addition to the AEDs. One study found that key challenges for parents were the disclosure of epilepsy and the lack of adequate information about coping in terms of psychosocial and emotional support.(20). Parents of children with chronic conditions require time to discuss their and their child’s issues and doctors may not be able to provide the amount of time required. Parents and children also require easy access to support services such as social workers and physiotherapists. One mother in our study suggested having a multidisciplinary epilepsy clinic including a paediatrician, social worker and physiotherapist. Paediatric nurses could run the clinic and a pharmacist could also be involved to assist with drug supply issues or complications of AEDs. This would seem to be an appropriate and practical model to work towards.

## Conclusions

Comprehensive care for children with epilepsy requires a good knowledge of the individual patient - their condition and comorbidities, their family, and their strengths and vulnerabilities. It requires following patients over a long time, and ready availability of services directed at the specific needs of the children and their families. The quality of life of a child with epilepsy is not purely the result of epilepsy itself but depends on the child’s resilience, parental and family well-being, family attitudes and social and cultural background.

## Acknowledgements

We thank the patients and their families for participating in the study.

